# Activation of the Integrated Stress Response in drug-tolerant melanoma cells confers vulnerability to mitoribosome-targeting antibiotics

**DOI:** 10.1101/2020.06.26.173492

**Authors:** Roberto Vendramin, Angelina Konnova, Sara Adnane, Sonia Cinque, Vicky Katopodi, Zorica Knezevic, Panangiotis Karras, Ewout Demesmaeker, Francesca M Bosisio, Lara Rizzotto, Oliver Bechter, Jean-Christophe Marine, Eleonora Leucci

## Abstract

Therapy resistance remains a major clinical challenge for the management of metastatic melanoma. Here we show that activation of the Integrated Stress Response (ISR), which we show is common in drug-tolerant and resistant melanoma, promotes selective synthesis of mitochondrial proteins in the cytosol. Since mitochondrial translation adapts to the influx of nuclear-encoded mitochondrial proteins, ISR activation indirectly enhances mitochondrial translation and makes these cells highly vulnerable to mitochondrial translation inhibitors. Treatment of melanoma with mitoribosome-targeting antibiotics, induces proteotoxic stress and significantly compromises the growth of *NRAS*-mutant and immunotherapy-resistant skin melanoma as well as uveal melanoma. Additionally, a triple BRAFi/MEKi/Tigecycline combination reduces intratumour heterogeneity by abrogating emergence of dedifferentiated drug-tolerant cells, and delayed or even prevented the development of resistance in BRAF^V600E^ PDX models. Consistently, a melanoma patient exposed to Doxycycline, a mitoribosome-targeting antibiotic commonly used to treat infections, experienced a complete and long-lasting response of a treatment-resistant lesion.

**Significance:** Our study indicates that the repurposing of mitoribosome-targeting antibiotics offers a rational salvage strategy for targeted therapy in *BRAF*-mutant melanoma, and a therapeutic option to target *NRAS*-driven and immunotherapy-resistant cutaneous melanoma and uveal melanomas.

## Introduction

Resistance to cancer therapy is a major limiting factor for the achievement of a cure. In metastatic melanoma, for instance, despite the recent breakthroughs in targeted therapy and Immune Checkpoint Blockade (ICB) (Larkin et al., 2015; Sosman et al., 2012), the 5-year survival rate still remains around 52%. Considering that drug discovery and development is a very slow and expensive process (Pushpakom et al., 2019), affected by high attrition rates (Waring et al., 2015), repurposing of existing de-risked compounds may offer an attractive alternative. However, a deep understanding of the mechanisms underlying drug resistance is essential before novel combinations can be rationally designed.

The most commonly accepted explanation for the inexorable development of therapy resistance invokes specific genetic alterations that are acquired by chance before or during treatment (Holohan et al., 2013). However, recent findings indicated that a subset of cancer cells are capable to survive the therapeutic insult by engaging specific transcriptional programs that confer them with drug-tolerant phenotypes (Sharma et al., 2010; Menon et al., 2015; Roesch et al., 2013; Su et al., 2017). These drug-tolerant cells (also known as persisters) provide a pool, commonly referred to as minimal residual disease (MRD), from which stable resistance is established. These findings indicated that eradicating these residual cells before stable resistance is acquired may offer new promising therapeutic avenues (Boumahdi and de Sauvage, 2019; Rambow et al., 2019). However, characterization of the cellular composition of MRD using single-cell approaches have recently highlighted that implementing such a strategy will come with its own challenges. Indeed, co-emergence -within the same MRD lesion- of four very distinct drug-tolerant states was observed following exposure of *BRAF*-mutant melanoma to MAPK-targeted therapy (Rambow et al., 2018). These included the Starved Melanoma Cell (SMC) state sharing transcriptomic features of nutrient-deprived cells (Rambow et al., 2018), a Neural Crest Stem-like Cell (NCSC) state, an invasive or mesenchymal-like state that was recently renamed undifferentiated state (Rambow et al., 2018; Tsoi et al., 2018) and a hyperdifferentiated state. The two de-differentiated states, NCSC and undifferentiated/mesenchymal, which harbour cancer stem cell features, are considered as particularly important drivers of tumour recurrence (Boshuizen et al., 2018; Rambow et al., 2018). Unfortunately, there are currently no clinically-compatible approaches known to efficiently co-target these two distinct cell populations. Whether these (and the other additional) drug-tolerant subpopulations although harbouring distinct gene expression signature-exhibit common and actionable vulnerabilities has therefore become a key question.

Although the metabolic profile of cancer cells varies across patients, tumour types and subclones within a tumour, there is emerging evidence that mitochondrial bioenergetics, biosynthesis and signaling are required for tumorigenesis. Accordingly, several studies have begun to demonstrate that mitochondrial metabolism is potentially a fruitful arena for cancer therapy (Jagust et al., 2019; Weinberg and Chandel, 2015). Critically, as our understanding of the biology of MRD increases across multiple tumour types, it is becoming clear that persister cells exhibit a dependency on mitochondrial biology that is exacerbated, even higher than those of their drug-naïve counterparts (Davis et al., 2020; Jagust et al., 2019; Sharon et al., 2019). For instance, relapse-initiating cells in B-progenitor Acute Lymphoblastic Leukemia (B-ALL) are characterized by elevated levels of mitochondrial metabolism (Dobson et al., 2020). As in B-ALL, a slow cycling population of melanoma cells that emerges in cultures exposed to vemurafenib or cisplatin exhibits elevated oxidative phosphorylation and targeting mitochondrial respiration blocks their emergence and delayed drug resistance (Roesch et al., 2013). It remains unclear, however, whether metabolic reprogramming is the only underlying cause of this increased mitochondrial dependency and whether (all) drug-tolerant cells that co-emerge following BRAF and MEK co-inhibition are equally sensitive to mitochondria-targeting agents.

Drug-tolerant cells from multiple cancer types often exhibit elevated activation of the Integrated Stress Response (ISR) (Deng and Haynes, 2017) following drug exposure (Almanza et al., 2019; McConkey, 2017) and the role of the key mediator of the ISR, ATF4, in drug tolerance is increasingly recognized (Jewer et al., 2020; Rzymski et al., 2009).The ISR is an adaptive translation program, triggered by several intracellular and extracellular stressors, that converges on the phosphorylation and activation of the eukaryotic translation initiation factor 2 (eIF2) complex. Activation of this pathway, results in reduced global translation alongside the selective synthesis of proteins important for survival. When the stress become overwhelming however, the same pathway triggers apoptosis (Verheyden et al., 2019). Emerging evidence indicates that in mammals the ISR is essential to convey mitochondrial disfunctions to the nucleus through a process known as retrograde signalling (Quirós et al., 2017). Conversely, the effect of ISR activation on mitochondrial activities is less clear. The observation that the rate of mitochondrial translation adapts to the influx of nuclear-encoded mitochondrial proteins (Richter-Dennerlein et al., 2016) raised the possibility that drug-tolerant cells may also tune-down their mitochondrial translation rate. In contrast to this prediction, however, we show herein that activation of ISR in drug-tolerant cells promotes selective translation of a subset mRNA encoding for mitochondrial proteins, and thereby generates an anterograde signalling from the cytosol boosting mitochondrial translation. This mechanism may explain the exquisite sensitivity of persister cells to mitochondria targeting agents such as uncouplers, and importantly identify mitochondrial translation as one critical sensitive node.

Interestingly, specific antibiotics can be repurposed to inhibit mitochondrial protein synthesis. Indeed, just like bacteria from which they originate from, the mitochondrial translational machinery of cancer cells, and in particular their ribosomes, are inhibited by tetracyclines (Kalghatgi et al., 2013; Moullan et al., 2015). The antitumor efficacy of these antibiotics has already been demonstrated *in vitro* (Ahler et al., 2013) and *in vivo* in haematological malignancies (D’Andrea et al., 2016; M et al., 2018; Skrtić et al., 2011; Zhang et al., 2017) and clinical trials in AML and double hit lymphomas are currently ongoing (Reed et al., 2016). Critically, we show herein that targeting mitochondrial protein synthesis with antibiotics of the tetracycline family prevented emergence of most (3 out of 4) drug tolerant subpopulations and delayed and even prevented the development of resistance to MAPK inhibition in *BRAF*-mutant preclinical PDX models as well as in one patient. Finally, we also showed efficacy of this treatment on models derived from patients with limited therapeutic options such as uveal melanoma, *BRAF* wild-type skin melanoma, or that had developed resistance to targeted and immunotherapies.

Together these findings indicate that the mitochondrial translation targeting with antibiotics of the tetracyclines family should be exploited to rationally design anticancer therapeutic regimens that could rapidly be implemented into patients, given the wide-spread clinical use of such agents.

## Results

### Activation of the Integrated Stress Response (ISR) increases mitochondrial translation

Phenotype switching into an undifferentiated drug-tolerant state can be induced *in vitro* by activating the ISR, leading to an ATF4-dependent decrease in MITF expression (Falletta et al., 2017). Accordingly, exposure of drug-naïve melanoma cells to salubrinal, an ISR agonist, increased levels of ATF4 and caused a concomitant downregulation of MITF (Figure 1A).

**Figure 1.**
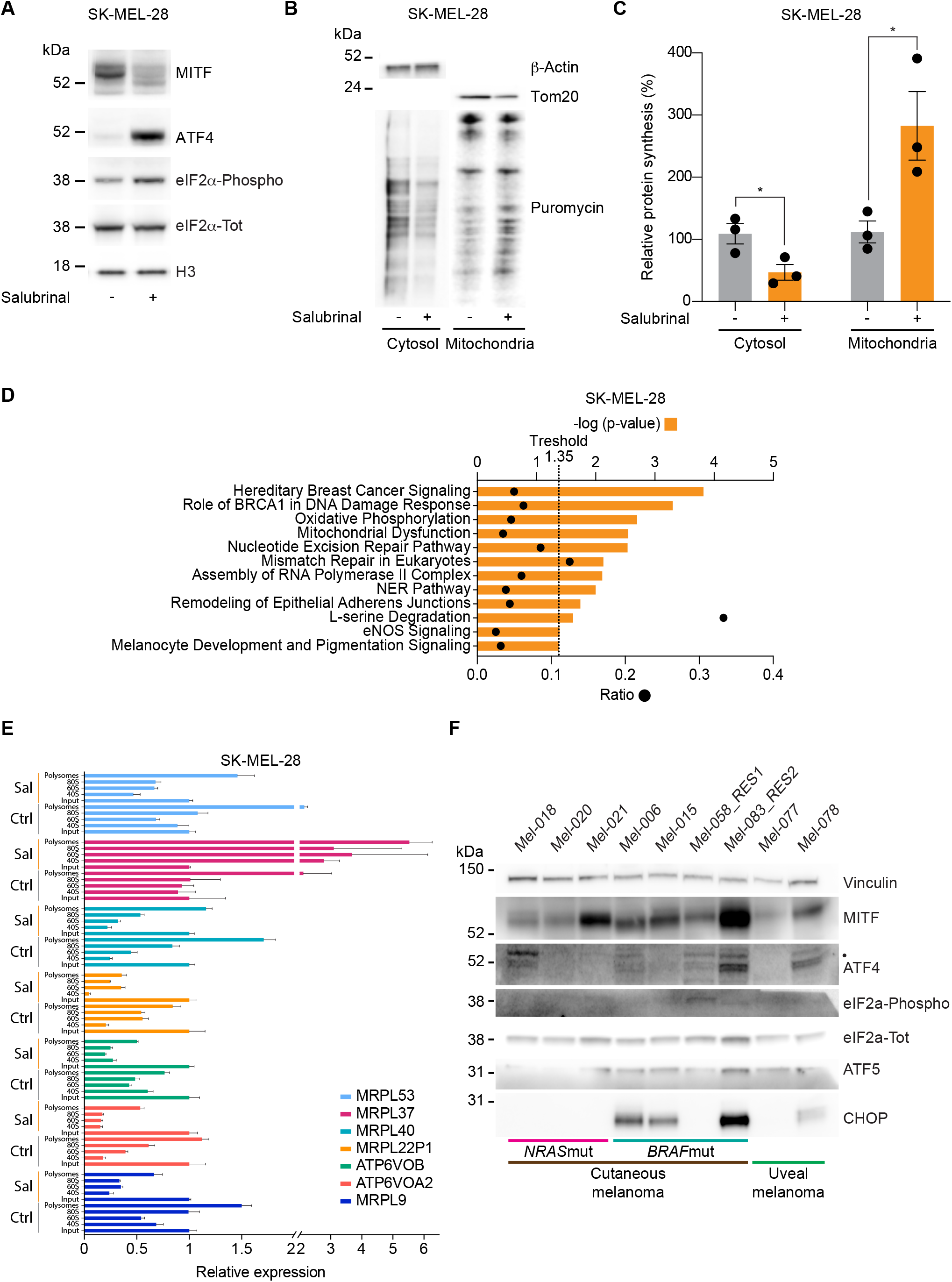
Activation of the Integrated Stress Response (ISR) increases the dependency of melanoma on mitochondrial translation. **A**, Western blotting of SK-MEL-28 cells 72 hours after treatment with salubrinal (+) or DMSO (-). Representative images of three independent experiments. **B,** Western blotting of cells described in **A**, after a 10-min pulse with puromycin and subsequent cytosol-mitochondria fractionation. Representative images of three independent experiments. **C,** Quantification of protein synthesis (%), measured by calculating the intensity of the puromycin signal on western blot, in SK-MEL-28 cells as described in **B**. Data are the means ± s.e.m. of different biological replicates. *P* value was calculated by Student’s t-test. **D,** Ingenuity Pathway Analysis (IPA) of most differentially expressed genes in RNA obtained from polysome profiling of cells described in **A**, with orange bars indicating −log (p-value) and dots indicating the ratios. **E,** RT-qPCR of cells described in D for mitochondrial encoded genes. Sal: Salubrinal, Ctrl: DMSO. Error bars represent mean ± S.D. of three independent experiments. **F,** Western blotting of a panel of drug-naïve melanoma PDX models. *\*P*<0.05. Mut: mutant; RES1: resistant to BRAFi. RES2: Resistant to BRAFi+MEKi and to anti-PD-1+anti-CTLA-4, • indicates the ATF4 band, mut: mutant.

The observation that drug-tolerant cells downregulate cytosolic protein synthesis suggests that these cells may also tune-down the activity of their mitochondrial translation machinery. Surprisingly however, puromycin incorporation assay followed by mitoplast isolation upon salubrinal treatment, demonstrated that ISR activation caused a dramatic increase in mitochondrial translation, and thus despite the expected overall decrease in cytosolic translation (Figures 1B and C). To further investigate the underlying mechanism, we identified translationally regulated mRNAs upon salubrinal treatment by performing polysome profiling analyses followed by RNA sequencing. We identified 382 transcripts whose association with ribosomes significantly (adj. p value<0.05) changed in response to the ISR activation and thus in response to phenotype switching and acquisition of therapy resistance. As expected, the vast majority of the transcripts (90%) were depleted from the ribosomal fraction upon ISR activation while only 10% showed an enrichment. Among those 2.3% where mitochondrial mRNAs. In line with this, Ingenuity Pathway Analysis shows enrichment in mitochondria-related terms (Figure 1D). These findings were further validated by performing polysome profiling followed by RT-qPCR upon salubrinal treatment (Figure 1E). Since the influx of nuclear-encoded mitochondrial proteins from the cytosol directly regulates mitochondrial translation (Richter-Dennerlein et al., 2016), the latter observation provides a likely explanation for the observed increased in mitochondrial protein synthesis. These observations also provide insights into the molecular mechanisms underlying increased dependency on mitochondrial metabolism, and identify mitochondrial translation as a putative Achiles’ heel of cells engaging the ISR pathway, such as drug-tolerant cells. Note that western blot analysis detected elevated levels of key ISR markers, such as P-eIF2α, ATF4 and CHOP, in non-drug-exposed human melanoma lesions, including lesions that were sensitive and resistant to both targeted and immune-therapies. This observation indicates that ISR signalling is not only activated in response to drug treatment (Figure 1F).

### Tetracyclines restrain the growth of therapy-resistant melanoma

We reasoned that the increasing dependency on mitochondrial translation during the development of therapy resistance should sensitise melanoma cells to antibiotic-induced mitochondrial translation block. To test this hypothesis, we exposed several cutaneous and *BRAF* wt (wildtype), uveal melanoma and immunotherapy-resistant IGR37 and YUMM 1.7 cell lines (together with its sensitive counterpart YUMMER 1.7; see Table1) to increasing concentrations of two different antibiotics: Tigecycline or Doxycycline (Figures 2A and B and Figures S1A and B). All cells lines showed a significant and dose-dependent decrease in cell growth with both antibiotics. We therefore tested the efficacy of antibiotics that target mito-ribosomes on the growth of several PDX melanoma models resistant to various therapeutic modalities (Figures 2C-F). A daily intraperitoneal (i.p.) injection of Tigecycline (50 mg/kg) was sufficient to effectively increase overall survival in NRAS^Q61R^-mutant (Figure 2C and Figure S2A). Milder, but significant, results were obtained in drug-naïve BRAF^V600E^-mutant cutaneous melanoma models sensitive to MAPKi (Figures S2B and C). Likewise, daily monotreatment of C57BL/6 mice engrafted with the immunotherapy-resistant YUMM 1.7 cells (Figure S2D) were sensitive to Tigecycline (Figure 2D). Furthermore, Tigecycline (50 mg/kg) treatment of a uveal melanoma PDX model Mel-077, derived from a patient progressing on the immune checkpoint inhibitor pembrolizumab (an anti-PD1 agent), was also sufficient to stabilise tumour growth and significantly delay progression (Figure 2E). Lastly, the immunotherapy-sensitive YUMMER 1.7 cells (derived from YUMM 1.7, (Wang et al., 2017)) were engrafted in C57BL/6 mice and treated with anti-PD1 alone or in combination with Tigecycline (50 mg/kg) to check whether the antibiotic treatment synergized with ICB. In line with previous findings in mice and humans, the anti-tumour effect of anti-PD1 alone was only measurable after a lag period of a few days (Figure 2F and Figure S2E). Adding Tigecycline to anti-PD1 resulted in a faster and more efficient tumour debulking (Figure 2F and Figure S2E). Together, these data indicated that exposure to tetracyclines alone is sufficient to significantly delay progression of melanoma lesions that are insensitive to targeted- and immune-therapies *in vivo*, and to accelerate anti-tumour immune responses when combined with ICB.

**Table 1.** Cell lines used in the study. Mutational and phenotypic status of cell lines used.

| **Cell line** | **Mutation** | **Phenotype** |
| --- | --- | --- |
| MM011 | NRAS<sup>Q61</sup> | Proliferative |
| MM099 | BRAF<sup>V600E</sup> | Invasive |
| MM165 | NRAS<sup>Q61</sup> | Invasive |
| WM383 | BRAF<sup>V600E</sup> | NCSC |
| WM852 | NRAS<sup>Q61</sup> | NCSC |
| IGR37 | BRAF<sup>V600E</sup> | Immunotherapy-resistant |
| YUMM 1.7 | BRAF<sup>V600E</sup>/wt PTEN<sup>-/-</sup> CDKN2<sup>-/-</sup> | Immunotherapy-resistant |
| YUMMER 1.7 | BRAF<sup>V600E</sup>/wt PTEN<sup>-/-</sup> CDKN2<sup>-/-</sup> | Immunotherapy -sensitive |
| UM 92.1 | GNAQ |  |

**Figure 2.**
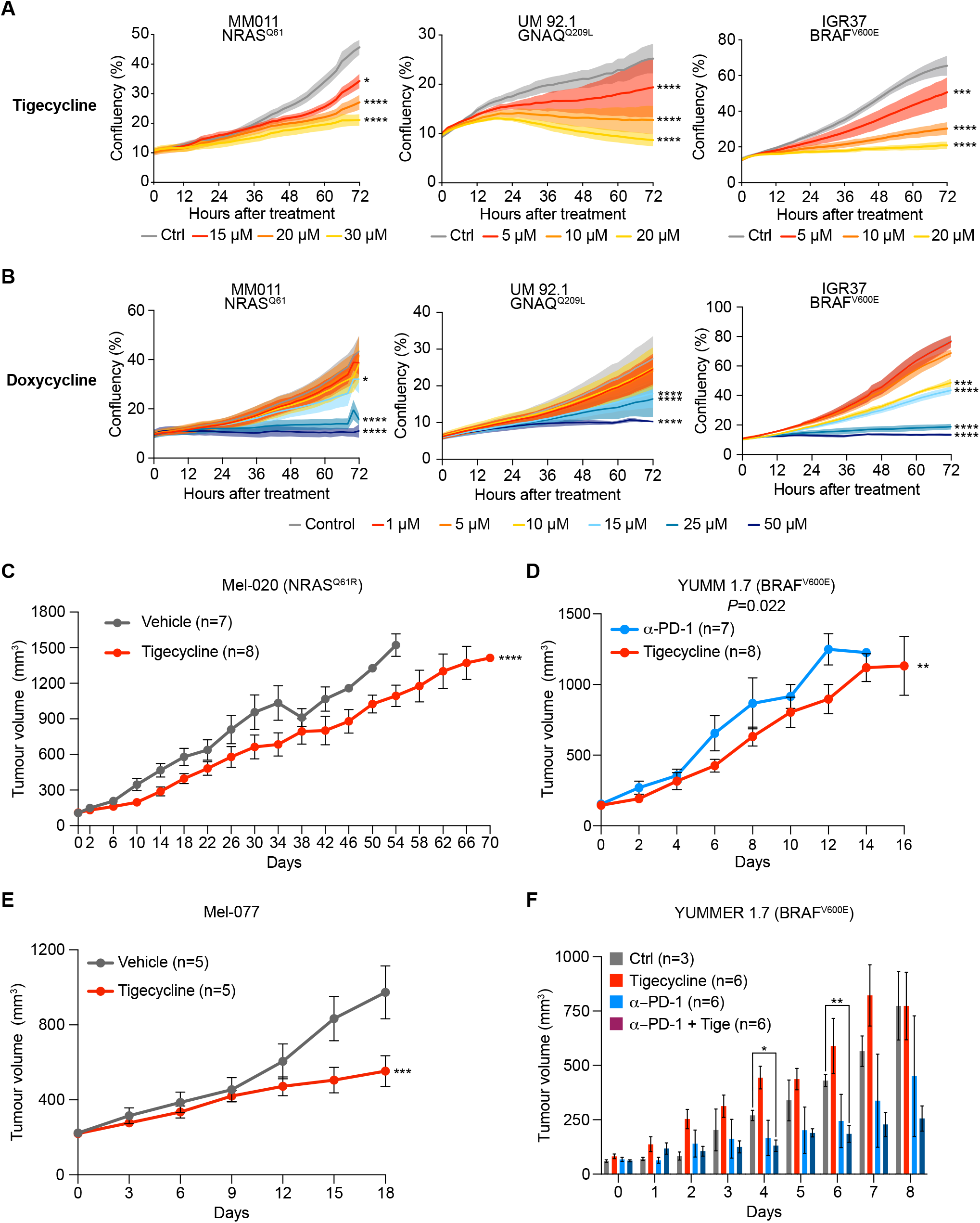
Tetracyclines restrain the growth of therapy-resistant melanomas. **A**, Cell growth (measured as % of cell confluency) of MM011 (*NRAS*mutant), UM 92.1 (GNAQ mutant, uveal melanoma) and IGR37 (resistant to immunotherapy) cell lines upon exposure to increasing concentrations of Tigecycline for 72 hours. Data are the means ± s.e.m. of three independent experiments. *P* values were calculated by Dunnett’s test. **B,** Cell growth (measured as % of cell confluency) of cells described in **A** upon exposure to increasing concentrations of Doxycycline for 72 hours. Data are the means ± s.e.m. of three independent experiments. *P* values were calculated by Dunnett’s test. **C,** Tumour volume of cohorts of Mel-020 NRAS^Q61R^ PDX mice treated with vehicle (DMSO) or Tigecycline. Data are the means ± s.e.m. of different biological replicates. *P* value was calculated by two-way ANOVA. **D,** Tumour volume of cohorts of YUMM 1.7 mouse xenografts treated with α-PD-1 or Tigecycline. Data are the means ± s.e.m. of different biological replicates. *P* value was calculated by two-way ANOVA. **E,** Tumour volume of cohorts of Mel-077 uveal melanoma PDX mice treated with vehicle (DMSO) or Tigecycline. Data are the means ± s.e.m of different biological replicates. *P* value was calculated by two-way ANOVA. **F,** Tumour volume of cohorts of YUMMER 1.7 mouse xenografts treated with α-PD-1, Tigecycline (Tige), a combination of the two or untreated (Ctrl). Data are the means ± s.e.m. of different biological replicates. *P* value was calculated by two-way ANOVA with Geisser-Greenhouse correction. *\*P*<0.05, *\*\*P*<0.01, \*\*\**P*<0.001, \*\*\*\**P*<0.0001.

### Tigecycline overcomes acquired resistance to MAPK inhibitors *in vivo* and significantly increases overall and progression-free survival

We next exposed several human melanoma cell lines (see Table1) that harbour transcriptional profiles reminiscent to two critical drug-tolerant states, namely invasive/undifferentiated and NCSCs, to increasing concentrations of both Tigecycline or Doxycycline. All established lines showed a significant and dose-dependent decrease in cell growth with both antibiotics (Figures 3A-C and Figure S3A). Moreover, western blot for ISR regulators in these cell lines confirmed that the ISR pathway is already activated before treatment and that exposure to the antibiotics alone or in combination with MAPK inhibitors (MAPKi) further exacerbated proteotoxic stress and triggered CHOP activation, which culminated in induction of cell death (Figure 3D and Figure S3B).

**Figure 3.**
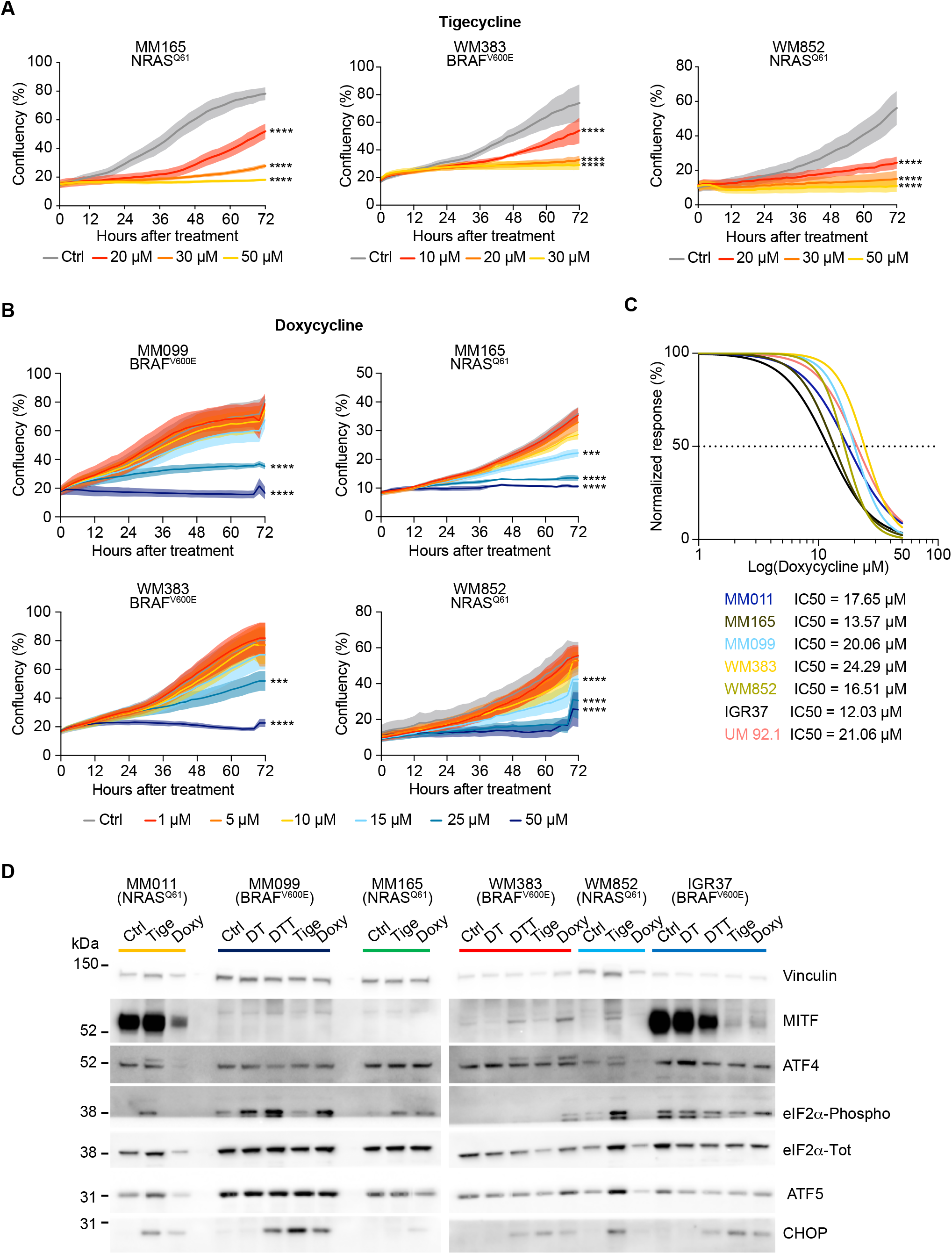
Tetracyclines exacerbate the activation of the ISR and affect the viability of multiple drug-tolerant states. **A,** Cell growth (measured as % of cell confluency) of MM165 (*NRAS*mutant, invasive), WM383 (*BRAF*mut, NCSC-like) and WM852 (*NRAS*mutant, NCSC-like) cell lines upon exposure to increasing concentrations of Tigecycline for 72 hours. Data are the means ± s.e.m. of three independent experiments. *P* values were calculated by Dunnett’s test. **B,** Cell growth (measured as % of cell confluency) of MM099 (*BRAF*mut, invasive) MM165 (*NRAS*mutant, invasive), WM383 (*BRAF*mut, NCSC-like) and WM852 (*NRAS*mutant, NCSC-like) cell lines upon exposure to increasing concentrations of Doxycycline for 72 hours. Data are the means ± s.e.m. of three independent experiments. *P* values were calculated by Dunnett’s test. **C,** Growth inhibition curves and IC50 values of several cutaneous melanoma cell lines and one uveal melanoma (UM 92.1) cell line upon exposure to increasing concentrations of Doxycycline. Data are the means ± s.e.m. of three independent experiments. **D,** Western blotting of a panel of different melanoma cell lines treated with vehicle (DMSO), Dabrafenib+Trametinib (DT), Dabrafenib+Trametinib+ Tigecycline (DTT), with Tigecycline (Tige) or with Doxycycline (Doxy) for 72 hours. Representative image of three independent experiments. *** *P*<0.001, \*\*\*\**P*<0.0001.

These observations indicated that tetracyclines may eradicate the residual tumour cells emerging following exposure to MAPK inhibition. We therefore tested whether addition of Tigecycline (T) to Dabrafenib and Trametinib (DT), a standard-of-care treatment for patients with *BRAF*-mutant melanoma, delayed or prevented the onset of resistance in two different BRAF^V600E^ melanoma PDXs (Mel-006 and Mel-015) (Figure 4). As expected, all mice from both cohorts initially responded to DT but, eventually, developed resistance (Figures 4A-D and Figures S4A and B).

**Figure 4.**
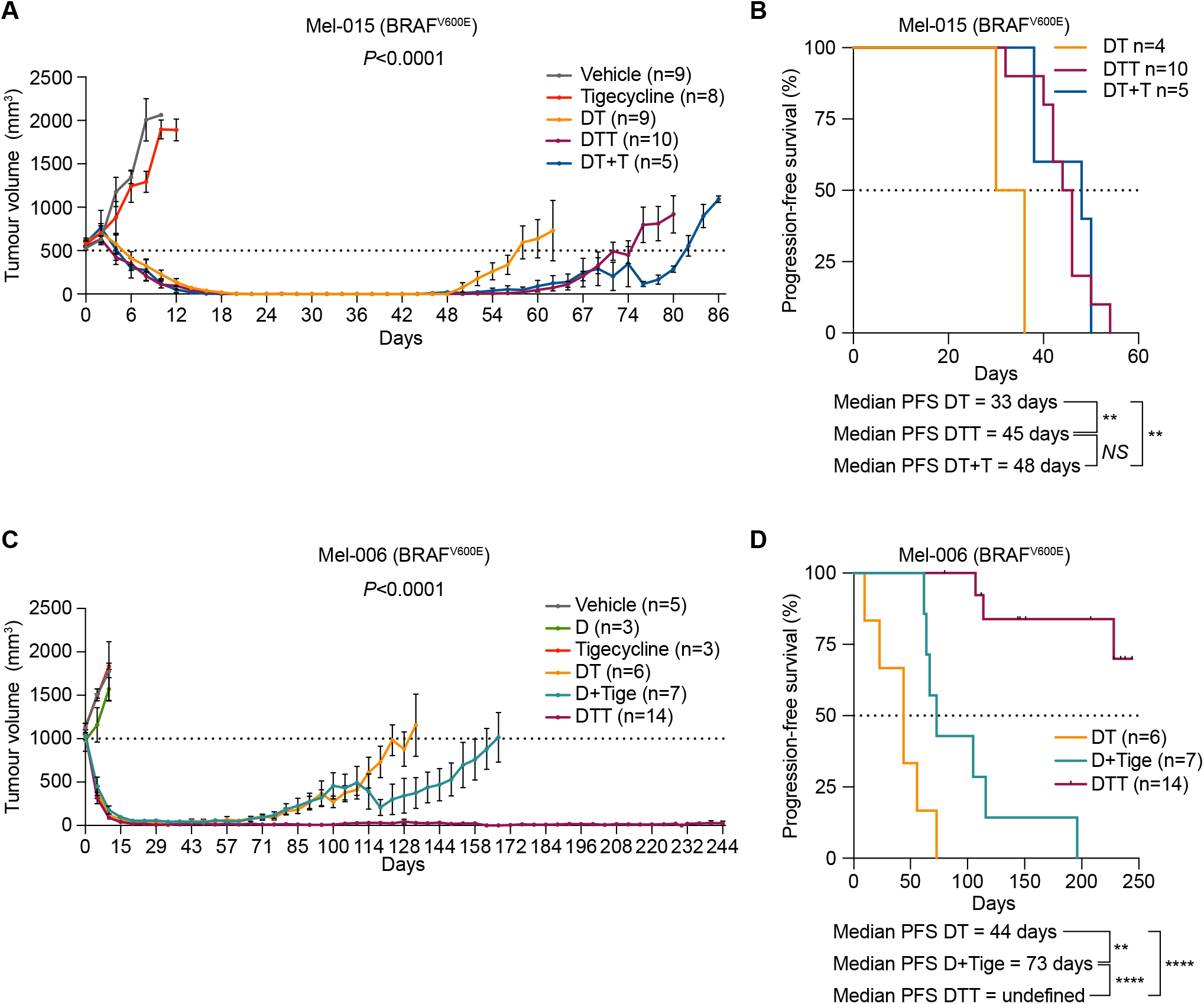
Tigecycline overcomes acquired resistance to MAPK inhibitors *in vivo* and significantly increases progression-free survival. **A,** Tumour volume of cohorts of Mel-015 BRAF^V600^ PDX mice treated with vehicle (DMSO), Tigecycline, Dabrafenib+Tigecycline (D+Tigecycline), Dabrafenib+Trametinib (DT), Dabrafenib+Trametinib+Tigecycline (DTT) or with Dabrafenib+Trametinib with the addition of Tigecycline at MRD (DT+T). Data are the means ± s.e.m. of different biological replicates. *P* value was calculated by two-way ANOVA. **B,** Kaplan-Meier-plot showing progression-free survival of mice described in (A). *P* values were calculated by log-rank (Mantel-Cox) test. **C,** Tumour volume of cohorts of Mel-006 BRAF^V600E^ PDX mice treated with vehicle (DMSO), Dabrafenib (D), Tigecycline, Dabrafenib+Trametinib (DT), Dabrafenib+Tigecycline (D+Tigecycline) or with Dabrafenib+Trametinib+Tigecycline (DTT). Data are the means ± s.e.m. of different biological replicates. *P* value was calculated by two-way ANOVA. **D,** Kaplan-Meier-plot showing progression-free survival of mice described in (C). *P* values were calculated by log-rank (Mantel-Cox) test. *NS: P*>0.05, *\*\*P*<0.01, \*\*\*\**P*<0.0001.

Notably, in the Mel-015 cohort (Figures 4A and B and Figure S4A) the addition of antibiotics, whether added from the beginning of the treatment (DTT) or after lesions reached MRD (DT+T), significant delayed the development of resistance. Strikingly in the Mel-006 cohort, the DTT treatment resulted in complete remission in 11 out of 14 mice. Importantly, complete remission was observed even when the DTT treatment was completely interrupted during the MRD phase (Figures 4C and D and Figure S4B). Notably, a double Dabrafenib plus Tigecycline combination (D+Tige) was as effective as a DT treatment at inducing tumour regression in this particular PDX model. In addition, it significantly increased progression-free survival (PFS) and overall survival (OS) compared to DT (Figures 4C and D and Figure S4B). Thus, addition of Tigecycline successfully delayed and/or prevented acquisition of resistance to targeted therapy in *BRAF*-mutant skin melanoma preclinical models.

### Tigecycline efficiently reduces intra-MRD heterogeneity

We have so far only provided evidence that the tetracyclines affect the growth of both mesenchymal and NCSC de-differentiated drug-tolerant cells only *in vitro*. To assess the sensitivity of these and other drug-tolerant subpopulations in an *in vivo* context, we treated Mel-006 either with a Dabrafenib plus Tigecycline combination (D+Tige) or with DT until reaching MRD. Compared to DT, the Dabrafenib plus Tigecycline combination successfully reduced intratumour heterogeneity by eradicating most of the NCSCs (NGFR^+^AQP1^+^), undifferentiated/mesenchymal-like cells (AXL+) and SMCs (CD36^+^) drug-tolerant cells (Figure 5A).

**Figure 5.**
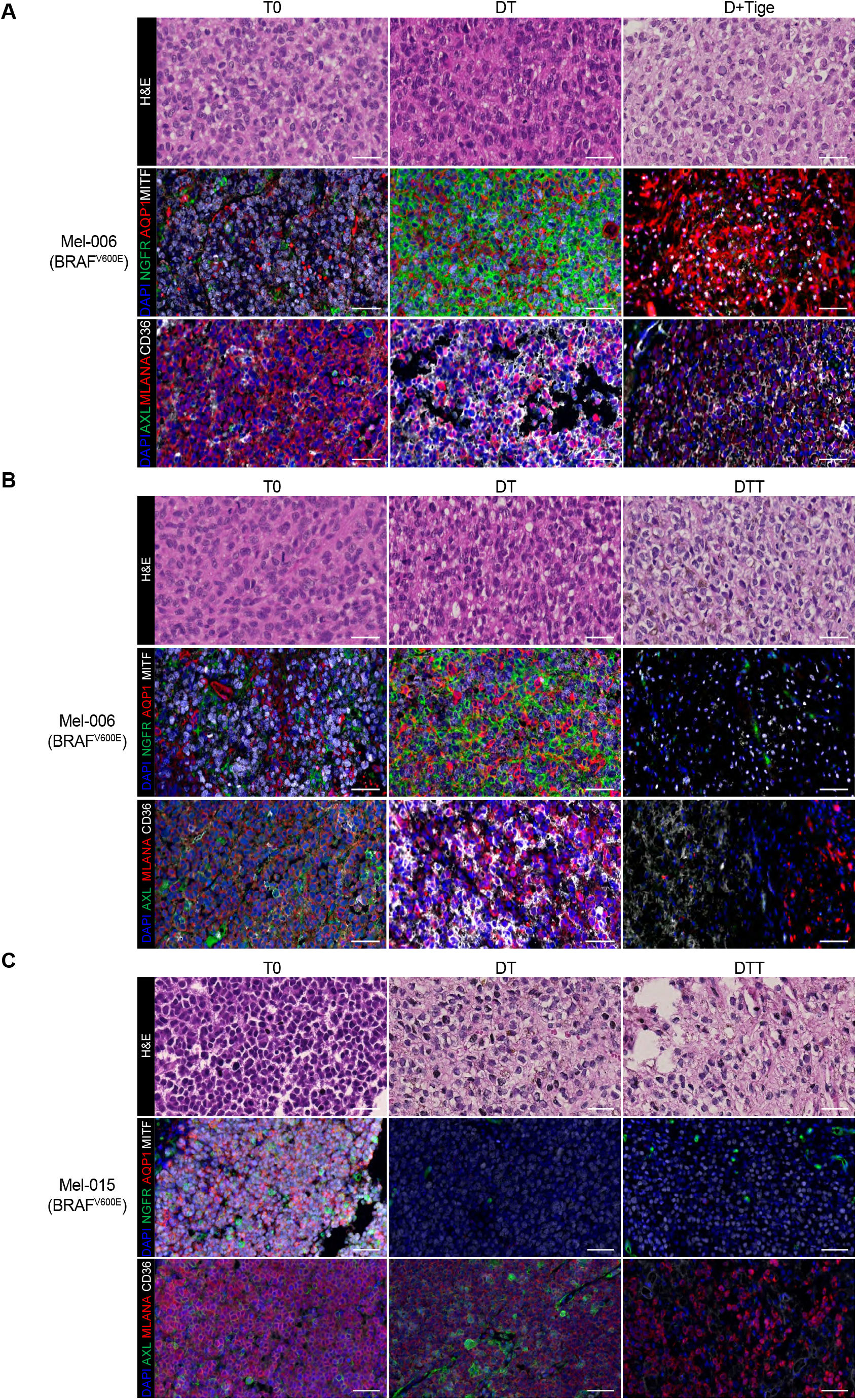
Tigecycline efficiently reduces intratumour heterogeneity. **A,** *Upper panel*, representative H&E staining of Mel-006 PDX tumours before treatment (T0, *left image*), on treatment with MAPK inhibitors at Minimal Residual Disease (MRD) (DT, *middle image*) and on treatment with Dabrafenib and Tigecycline (D+Tige, *right image*) at MRD. *Middle panel*, representative immunofluorescence staining against MITF (white), NGFR (green) and AQP1 (red) before treatment (T0) and at MRD of DT and D-Tige lesions. *Lower panel*, representative immunofluorescence staining against CD36 (white), AXL (green) and MLANA (red) before treatment (T0) and at MRD of DT and D-Tige lesions. Slides were counterstained with DAPI (blue). Scalebar = 50 μm. **B,** *Upper panel*, representative H&E staining of Mel-006 PDX tumours before treatment (T0, *left image*), on treatment at MRD with MAPK inhibitors (DT, *middle image*) and on treatment with Dabrafenib/Trametinib and Tigecycline (DTT, *right image*). *Middle panel*, representative immunofluorescence staining against MITF (white), NGFR (green) and AQP1 (red) before treatment (T0) and at MRD of DT and DTT lesions. *Lower panel*, representative immunofluorescence staining against CD36 (white), AXL (green) and MLANA (red) before treatment (T0) and at MRD of DT and DTT lesions. Slides were counterstained with DAPI (blue). Scalebar = 50 μm. **C,** *Upper panel*, representative H&E staining of Mel-015 PDX tumours before treatment (T0, *left image*), on treatment at MRD with MAPK inhibitors (DT, *middle image*) and on treatment with Dabrafenib/Trametinib and Tigecycline (DTT, *right image*). *Middle panel*, representative immunofluorescence staining against MITF (white), NGFR (green) and AQP1 (red) before treatment (T0) and at MRD of DT and DTT lesions. *Lower panel*, representative immunofluorescence staining against CD36 (white), AXL (green) and MLANA (red) before treatment (T0) and at MRD of DT and DTT lesions. Slides were counterstained with DAPI (blue). Scalebar = 50 μm.

Similarly, in two different *BRAF*-mutant PDX cohorts (Mel-006 and Mel-015) treated with the triple DTT combination until reaching MRD, the antibiotic successfully eradicated the NCSCs (NGFR^+^AQP1^+^), undifferentiated/mesenchymal-like cells (AXL^+^) and pseudo-starved cell population (CD36^+^). The hyperpigmented cell population (MITF^+^MLANA^+^) was the only drug-tolerant population to resist the combinatorial treatment (Figures 5B and C). These data indicated that a combination of antibiotics and MAPK inhibitors reduces intra-MRD heterogeneity by targeting mostly both de-differentiated (NCSC) and undifferentiated/invasive states.

### Activation of the ISR predicts durable responses to antibiotic treatment

We postulated that the difference in the efficacy of the antibiotic combinatorial treatment in preventing the development of resistance between the Mel-006 and Mel-015 BRAF^V600E^-mutant melanoma PDXs may result from a difference in the kinetic and/or extent at which the ISR pathway is activated. Indeed, in the Mel-006 PDX model, ATF4 activation was readily detectable before the start of the therapy (T0), an observation that may explain the high sensitivity to the antibiotic treatment and the long-lasting response (Figure 1D and 6A). In contrast, in the Mel-015 PDX model, ATF4 activation was only detected ON therapy (Figure 6B). Note that similar results were obtained in the Mel-077 UM model and in the Mel-020 NRAS^Q61R^-mutant model (Figures S5A and B). This observation may explain why the overall and progression-free survival was comparable in cohorts exposed to the antibiotics at MRD or from the beginning of the treatment. These data suggest that activation of apoptosis is likely to be a consequence of chronic activation of the ISR. In keeping with this, co-treatment with ISRIB, an inhibitor that blocks signaling downstream of the eIF2α kinase and consequently ATF4 activation, rescued apoptotic cell death (Figure 6C). These data underline that dependency on mitochondrial translation is a direct effect of ATF4 activation and that elevation of ATF4 levels is a predictive factor for durable response to antibiotic treatment.

**Figure 6.**
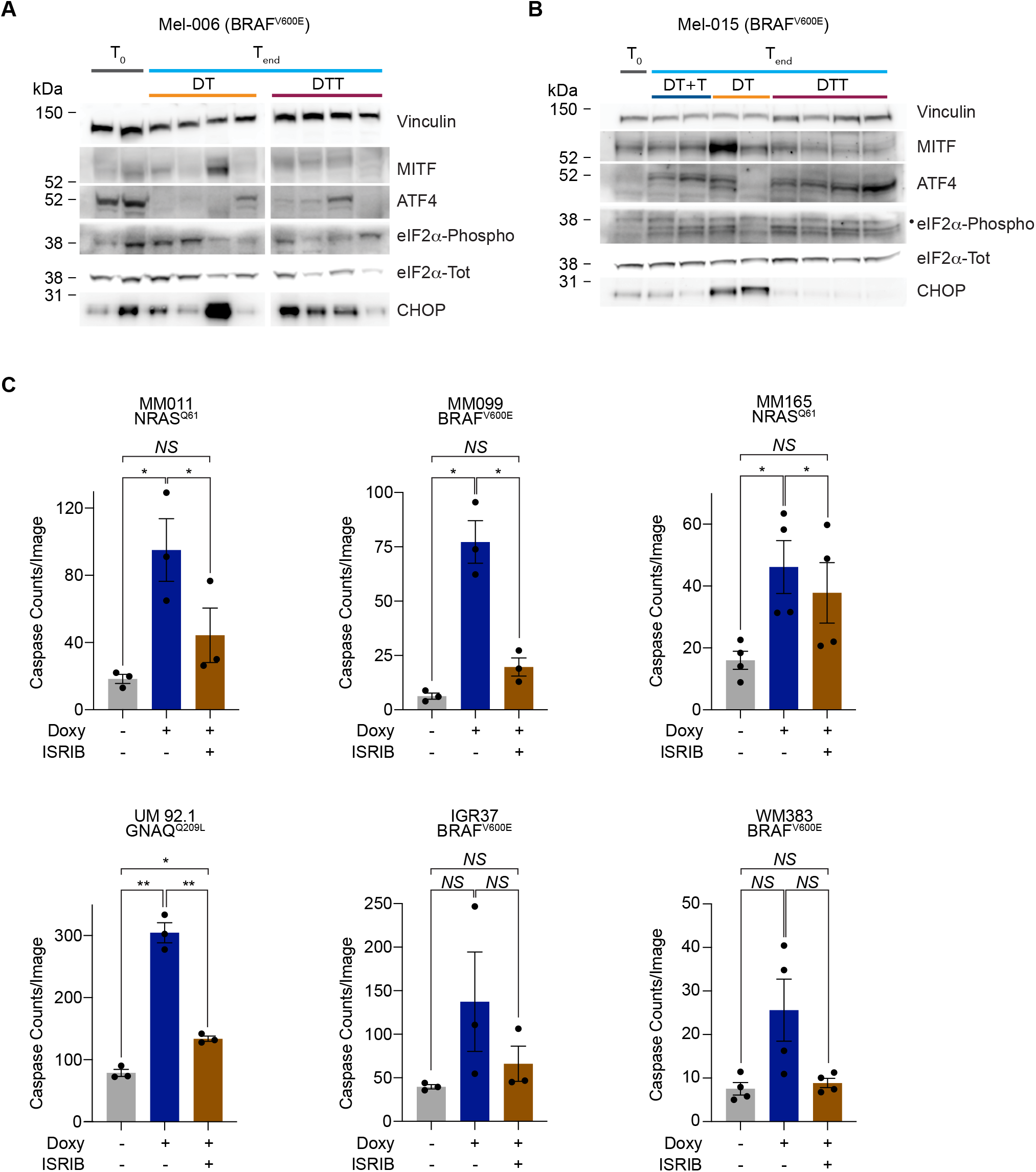
Activation of the ISR predicts durable responses to antibiotic treatment. **A,** Western blotting of Mel-006 (*BRAF*mut) PDX tumours collected right before the start of the treatment (T0) or at the end of the experiment (Tend) after receiving a daily dose of Dabrafenib+Trametinib (DT), or Dabrafenib+Trametinib+Tigecycline (DTT). **B,** Western blotting of Mel-015 (*BRAF*mut) PDX tumours collected right before the start of the treatment (T0) or at the end of the experiment (Tend) after receiving a daily dose of Dabrafenib+Trametinib (DT), Dabrafenib+Trametinib+Tigecycline (DTT) or with Dabrafenib+Trametinib with the addition of Tigecycline at MRD (DT+T). **C,** Caspase counts per image in a panel of cell lines treated with Doxycycline (Doxy, 50 μM) ± ISRIB (200 nM) for 72 hours. Data are the means ± s.e.m. of different biological replicates. *P* value was calculated by Student’s t-test. *NS: P*>0.05, *\*P*<0.05. • indicates the eIF2α-Phospho band.

### Doxycycline sensitises a MAPKi-refractory lesion in *BRAF*-mutant patients

Doxycycline is a broad-spectrum antibiotic belonging to the tetracycline family, which exerts its antibacterial action by binding to the 30S ribosomal subunit, thereby blocking translation. It can be administered orally for extended periods of time (Tan et al., 2011) with only minor adverse effects in patients. Importantly, it is often use in cancer patients under therapy when infections arise.

To assess the relevance of our *in vitro* and preclinical findings, we followed the clinical course of a 73-year-old female patient at our clinic. The patient was diagnosed with stage III melanoma in 2011 and with a relapse of her disease in 2016 after an episode of jaundice which was related to a tumour mass in her gallbladder and bile duct. Biopsy of the mass showed malignant melanoma (Figure 7A). Baseline PET CT (Figure 7B) showed metastatic tumour at the aforementioned locations as well as in the liver.

**Figure 7.**
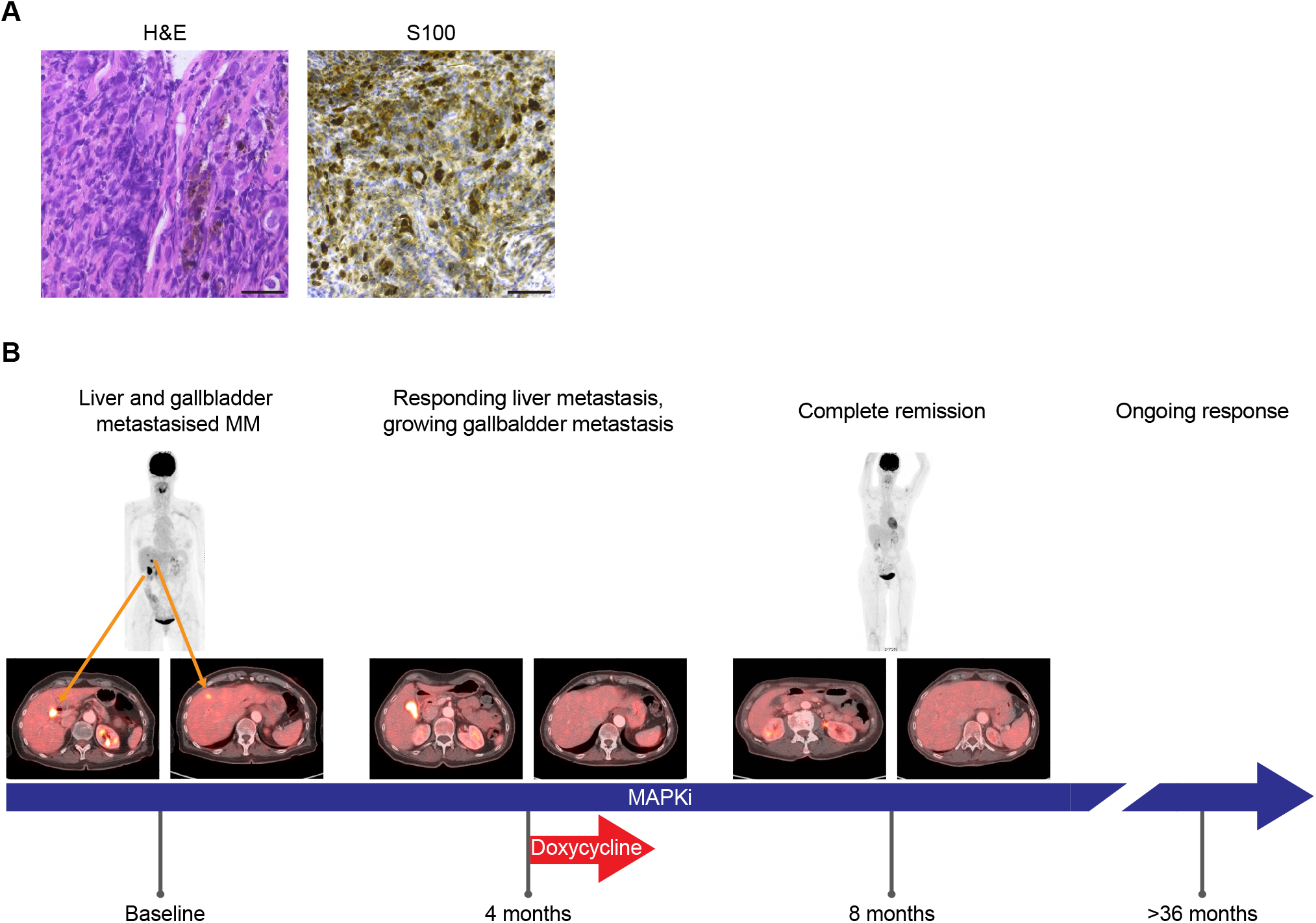
Doxycycline sensitises melanoma to MAPKi in patients. **A,** Histology and immunohistochemistry of the bile duct biopsy confirming the presence of melanoma metastasis. Scalebar = 25 μm. **B,** Case presentation of a patient with MAPK inhibitory therapy and sporadic exposure to Doxycycline. Baseline PET-CT revealed liver (*right lower-left image*) and gallbladder metastasis (*left lower-left image*). First response assessment showed disappearance of the liver metastasis but a persistent gallbladder lesion (4 months of treatment). Adding Doxycycline to MAPK inhibitory therapy showed shrinkage after two more cycles of treatment (CT scan not shown) which was subsequently confirmed by a PET-CT after 8 months on treatment revealing a complete metabolic response. After >36 months the patient is still responding to the treatment.

Subsequently BRAF-MEK inhibitor therapy was started. Response assessment after 4 cycles showed that while the liver metastasis had responded to the treatment, the gallbladder did not. Due to a skin infection the patient received Doxycycline during her 5^th^ BRAF-MEK inhibitor cycle for a total of 12 days. After 6 cycles the gallbladder metastasis started to regress, as measured by a CT scan, and the PET-CT scan after 8 cycles showed a complete response. The patient is still under BRAF-MEK inhibitor therapy and has a persistent response now lasting for more than 36 months (Figure 7A). This very unique case study provides proof-of-concept evidence that supports the sensitizing effect of tetracyclines to MAPK inhibitors in a clinical setting.

## Discussion

An emerging theme in cancer biology and drug resistance is the reliance of different lesions and drug tolerant states, across multiple cancer types, on mitochondria (Chen, 2012; Davis et al., 2020; Faubert et al., 2020; Haq et al., 2013; Jagust et al., 2019). The exact mechanism undelaying dependency of metabolically distinct lesions on mitochondrial however, remains unclear. Here we report that activation of the ISR, which is common to multiple drug-tolerant cells, promotes selective translation of mitochondrial mRNAs in the cytosol. As a result, melanoma cells increase mitochondrial translation, and thereby their vulnerability to inhibitors of mitochondrial protein synthesis. Although the importance of the ISR for the retrograde signalling of mitochondrial disfunctions is emerging (Fessler et al., 2020; Guo et al., 2020; Molenaars et al., 2020), its role in promoting mitochondrial translation through anterograde signalling is novel and unexpected and underlines the importance of cytosolic adaptive responses in the generation of therapy resistance.

Accordingly, we show that tetracyclines addition to the standard-of-care targeted therapy (*i.e.* DT) extended PFS and even completely abolished relapse in a *BRAF*-mutant preclinical PDX model (Mel-006). Mechanistically, we showed that the antibiotics compromised the emergence and/or survival of multiple drug-tolerant subpopulations, including the NCSC, mesenchymal/undifferentiated and pseudo-starved subpopulations at MRD. Importantly, de-differentiation into the invasive/mesenchymal and NCSC states was reported as an escape mechanism to various anti-melanoma therapies, including T-cell transfer therapy (Landsberg et al., 2012; Lee et al., 2020; Mehta et al., 2018; Rambow et al., 2018).

Although still significantly delaying disease progression, the triple DTT combination did not prevent relapse in another *BRAF*-mutant preclinical PDX model (Mel-015). Importantly, we showed that the exacerbated response to the treatment was correlated with the level of ISR activation at steady state, before treatment. Levels of ATF4 were readily detectable in Mel-006, but not Mel-015, drug-naïve lesions. Equally interestingly, detectable levels of ATF4 predicted response of therapy-resistant melanoma to tetracyclines alone, such as uveal melanoma and *BRAF* wt cutaneous melanoma. These observations offer a new avenue for the treatment of these diseases with no, or only limited, therapeutic options and identify ATF4 levels as a predictive marker of long-lasting responses to tetracyclines used either alone or in combination with standard-of-care.

Moreover, there is increasing evidence that primary resistance to immunotherapy may also be driven by the upregulation of mitochondrial translation (Jerby-Arnon et al., 2018; Poźniak et al., 2019). In keeping with this, exposure to tigecycline accelerated the response to ICB in syngeneic models. These observations warrant the implementation into the clinic of a sequential treatment regimen in which tumour debulking is induced by the DTT combination followed by an immunotherapy approach aiming at eradicating the residual pool of hyperdifferentiated cells.

Recent reports highlighted that brain metastasis across different cancers (Chen, 2012; Chen et al., 2007; Cheng et al., 2019), including melanoma (Fischer et al., 2019; Sundstrøm et al., 2019), display increased oxygen consumption and are therefore more dependent on OXPHOS. Considering that tetracycline family antibiotics can cross the blood brain barrier (Nau et al., 2010), this approach may also be a viable option for patients with this life-threatening condition.

In addition to their ability to inhibit mitochondrial translation, it remains possible that the antitumor effect of tetracyclines is multifactorial. Tetracyclines are known to induce a glycolytic phenotype, in addition to the effect on mitochondrial ribosomes (Ahler et al., 2013). This effect may also contribute to the enhanced sensitivity of tumour cells to BRAF/MEK inhibition. In addition, both dabrafenib and trametinib are partially metabolized by CYP3A4 (Lawrence et al., 2014), which is inhibited by Tetracycline. Its co-administration may therefore raise the active concentrations of dabrafenib and trametinib and thereby enhanced anti-tumour efficacy of DT (Bassi et al., 2004). We also cannot rule out that, in addition to cancer cell intrinsic effects, the dramatic responses observed in the pre- and clinical settings are a consequence of tetracyclines remodelling the tumour microenvironment, such as the innate immune system and/or (intratumor) microbiome. This may be important as the efficacy of anti-PD-1 therapy has been shown to be affected by the microbiome (Matson et al., 2018; McQuade et al., 2019). However, most common symbiotic bacteria are resistant to Doxycycline, which is a broad-spectrum antibiotic that can be administered orally for long periods of time with no or minor toxic effects (Tan et al., 2011). Its administration is therefore unlikely to affect significantly the microbiome of patients.

In conclusion, we show that while cancer cells have the ability to survive a wide range of insults including therapy exposure-through ISR-dependent translation remodelling, they indirectly enhance mitochondrial translation and thereby their vulnerability to inhibitor of mitoribosome assembly or function. Consequently, the repurposing of mitoribosome-targeting antibiotics offers a new promising therapeutic avenue for the treatment of a large spectrum of melanoma patients, including those with limited therapeutic options. Importantly, this combination strategy can be easily implemented and further evaluated in the clinic. Moreover, we provide evidence that patient stratification maybe guided by ATF4 levels, which could be used an accompanying biomarker to predict efficacy.

## Acknowledgments

The present study was funded by Kom op tegen Kanker (Stand up to Cancer), the Flemish cancer society. E.L. is funded by the Melanoma Research Alliance (MRA) Amanda and Jonathan Eilian young investigator award. Trace staff is supported by the Stichting Tegen Kanker grant 2016-054. V.K. is a recipient of the FWO PhD fellowship 1S47519N. S.C. is a recipient of the FWO PhD fellowship 1SD1620N.

The authors wish to thank Ghanem E. Ghanem (Jules Bordet Institute), Marcus Bosenberg (Yale University), Irwin Davidson (Institute of Genetic and Molecular and Cellular Biology), Aart Jochemsen (Leiden University) and Göran Jönsson (Lund University) for kindly gifting some of the melanoma cell lines used in this study. We would also like to thank Amanda Lund for comments and suggestions.

## Author contribution

A.K. and R.V., performed and designed all the *in vitro* and the allograft experiments, S.A. performed the uveal melanoma PDX experiment and contributed to the *in vitro* experiments of Figures 1F, 6 A and B and Figures S5A and B, S.C. and V.K. performed the *in vitro* experiments in Figure 1 A-E. P.K. and M.F.B. performed and interpreted multiplex IHC, E.D. and L.R. performed and designed cutaneous melanoma PDX experiments, O.B. provided and analysed the clinical case, E.L. and R.V. interpreted the data, J.C.M. and E.L. designed the research study and wrote the manuscript with the input from all the authors.

## Methods

### Study design

The objectives of this study were (I) to identify novel metastatic melanoma vulnerabilities, (II) to prevent or delay acquisition of therapy resistance, (III) to sensitise therapy-resistant lesions to therapy and (IV) to exploit this knowledge for therapeutic benefit in a preclinical setting. To test this objective, we made use of several clinically relevant mouse models of metastatic cutaneous and uveal melanoma with different mutational background and therapy sensitivity (*i.e.* with intrinsic or acquired therapy resistance profiles). Mice were evaluated for overall survival, progression-free survival, tumour growth on a daily basis. Cell lines were monitored for cell growth and cell death. The melanoma patient was monitored for tumour load and tumour response to the treatments. Each *in vitro* experiment was repeated at least three times (as specified in the figure legend) to ensure reproducibility. For *in vivo* experiments, sample size was calculated using the software G power version 3.1.9.2 with a power of 80% to 95% and an alfa-error probability of 0.05%, assuming an effect size of 0.25 (based on previous experiments and pilot studies). This number was not altered during the experiment. This study included several cell cultures, two animal models and one melanoma patient. Investigators were not blinded.

### Cell Lines

SK-MEL-28 cells, WM cell lines (a kind gift from Göran Jönsson) and the IGR37 cell line (a kind gift from Irwin Davidson) were grown in RPMI 1640 (Gibco, Invitrogen) supplemented with 10% FBS (Gibco, Invitrogen). The patient-derived low-passage MM cell lines (a gift from G.-E. Ghanem) were grown in F-10 (Gibco, Invitrogen), supplemented with 10% FBS (Gibco, Invitrogen). The YUMM 1.7 and YUMMER 1.7 cell lines (a kind gift from Marcus Bosemberg (Meeth et al., 2016; Wang et al., 2017)) were grown in DMEM/F12 (50:50, Gibco, Invitrogen) media supplemented with and 10% FBS (Gibco, Invitrogen). The UM92.1 cell line (a kind gift from Aart Jochemsen) were grown in a 1:1 mixture of RPMI 1640 (Gibco, Invitrogen) and DMEM/F12 (1:1 mixture, Gibco, Invitrogen) media supplemented with and 10% FBS (Gibco, Invitrogen).

All cell lines used are of human origin, save for the YUMM 1.7 and YUMMER 1.7 cell lines, derived from male C57BL/6 (*Mus Musculus*). Gender of the patients from whom the cell cultures were derived is as follows (female: F; male: M): SK-MEL-28: M; MM011: F; MM099: M; MM165: M, IGR37: M, YUMM 1.7: M, UM92.1: F. All cell lines were confirmed negative for Mycoplasma prior to the study using the MycoAlert Mycoplasma Detection Kit (Lonza) according to the manufacturer’s specifications.

### Cell viability assays

For colony formation assays, cells were plated in six-well plates at the appropriate density (1.5 10^4^/well in the case of the MM011, MM099, MM165, WM383, WM852, IGR37, YUMM 1.7 and YUMMER 1.7 and 8 10^4^/well in the case of the UM92.1 cells). Cells were then treated with increasing amounts of Tigecycline and cultured for 3 days (YUMM 1.7, YUMMER 1.7 and UM 92.1), 5 days (WM852 and IGR37) or 7 days (MM011, MM165 and WM383). Cells were then washed twice with PBS, fixed and stained for 15 min with a 1% crystal violet in 35% methanol solution. Surface occupancy was measured using ImageJ.

For IncuCyte Proliferation Assays, cells were plated in 96-well plates (TPP) at the appropriate density (MM011, MM165, MM099, IGR37 and WM383 1.5 x 10^4^/well, WM852 5 x 10^3^/well, YUMM 1.7, YUMMER 1.7 and UM 92.1 2.5 x 10^3^/well). Cells were treated with increasing amounts of Tigecycline or Doxycycline and cultured for 72 hours. Apoptotic cells were labelled with the IncuCyte Caspase 3/7 Green Apoptosis Assay Reagent (Essen BioScience). Four images per well were taken at 2-hour intervals using an IncuCyte ZOOM system (Essen BioScience). The percentage of cell confluency and fluorescent green counts indicating apoptotic cells were measured and analysed by the IncuCyte ZOOM software.

### Polysome profiling

SK-MEL-28 cells (5 15 cm ∅ plates per each condition) were plated in order to have 70% confluency after 96h. The following day, cells were treated with 20 μM Salubrinal (Sigma-Aldrich). 72 hours after the start of the treatment cells were treated with 100 μg mL^−1^ of Cycloheximide (CHX, Sigma-Aldrich) for 12 minutes at 37 °C, collected and resuspended in lysis buffer (30 mM Tris-HCl, 150 mM KCl, 10 mM MgCl2 supplemented with DDT (Sigma-Aldrich) 1 mM, CHX 100 μg mL^−1^, SUPERase-IN RNase Inhibitor 20 U/μl (Invitrogen, Thermo Fisher Scientific), 1× Halt Protease and Phosphatase Inhibitor Single-Use Cocktail (Life Technologies) before the start of the experiment). Lysates were then incubated agitating at 4 °C for 35 minutes, then centrifuged at 17.000 rcf for 15 minutes at 4 °C. Lysates were then loaded on a sucrose gradient (the linear sucrose gradient 5-20% was generated from 2 different solutions, sucrose 5% and 20% made with Buffer G (20 mM Tris-HCl, 100 mM KCl, 10 mM MgCl2 supplemented with DDT 1mM and CHX 100 μg mL^−1^ before the start of the experiment)). Samples were then centrifugated in an SW41Ti rotor (Beckman Coulter) at 37.000 rpm for 170 minutes at 4 °C. The fractions were obtained with a BioLogic LP System (Bio-Rad). 14 fractions were collected from each sample, with each fraction having a final volume of 600 μL. From the initial 14 fractions, 4 final samples were obtained (by pulling together some of them): 40S, 60S, 80S and polysomes.

### RNA extraction

For RNA extraction from the fractions, each faction was digested at 37 °C for 90 minutes in a digestion mix of Proteinase K (final concentration 100 μg mL^−1^, Sigma-Aldrich) and SDS 1%. Phenol acid chloroform 5:1 (Sigma-Aldrich) and NaCl 10 mM were then added. Samples were centrifuged at 16.000 rcf for 5 minutes at 4 °C. The upper aqueous phase was transferred in a new tube and 1 mL of isopropanol was added. Samples were stored at −80 °C overnight to precipitate the RNA. The following day the samples were centrifuged at 16.000 rcf for 40 minutes at 4 °C. The supernatant was discarded and the pellet washed with 500 μL of 70% EtOH, then centrifuged again at 16.000 rcf for 5 minutes at 4 °C. Pellet were air dried and resuspended in DEPC water. No DNase treatment was performed (not needed). Input RNAs were extracted with TRIzol (Invitrogen) according to the manufacturer’s instructions.

### RT-qPCR

RNA was reverse transcribed using the High-Capacity complementary DNA Reverse Transcription Kit (Thermo Fisher Scientific) on a Veriti 96-well thermal cycler (Thermo Fisher Scientific). Gene expression was measured by qPCR on a QuantStudio 5 (Thermo Fisher Scientific) and normalized in qbase+ 3.0 (Biogazelle) using 28S and 18S as reference genes. Sequences of the primers are indicated in Supplementary Table 1.

### RNA sequencing and data analysis

Samples were prepared for sequencing with TruSeq Stranded Total RNA kit (Illumina) according to the manufacturer’s instructions. Libraries were sequenced on Illumina NextSeq according to the manufacturer’s instructions generating PE HO cycles: (R1: 100) - (I1: 6) - (I2: 0) - (R2: 100). Quality control of raw reads was performed with FastQC v0.11.5. Adapters were filtered with ea-utils v1.2.2.18. Splice-aware alignment was performed with STAR against the human hg19. The number of allowed mismatches was 2. Reads that mapped to more than one site to the reference genome were discarded. The minimal score of alignment quality to be included in count analysis was 10. Resulting SAM and BAM alignment files were handled with Samtools v0.1.19.24. Quantification of reads per gene was performed with HT-Seq count v0.5.3p3. Count-based differential expression analysis was done with R-based (The R Foundation for Statistical Computing, Vienna, Austria) Bioconductor package DESeq. Reported *P*-values were adjusted for multiple testing with the Benjamini-Hochberg procedure, which controls false discovery rate (FDR). List of differentially expressed genes were selected at FDR 1.35.

### Puromycin incorporation assay (SUNsET)

SUNsET was performed as described in (Schmidt et al., 2009). Briefly, ≈80% confluent adherent cells were washed twice in 1× PBS and subsequently pulsed with puromycin-containing media (InvivoGen, 10 μg mL^−1^) for 10 min. The cells were then supplemented with normal media for 60 min before downstream applications (chase). Due to the fact that puromycin is a structural analogue of aminoacyl tRNAs, it gets incorporated into the nascent polypeptide chain and prevents elongation. Therefore, when used for reduced amounts of time, puromycin incorporation in neosynthesized proteins directly reflects the rate of mRNA translation in vitro. Puromycin incorporation was measured by western blotting using an antibody that recognises puromycin.

### Cellular fractionation and mitoplast isolation

Briefly, mitochondria were purified from 4-6 × 10^7^ cells using mitochondria isolation kit for cultured cells (Thermo Fisher Scientific) according to manufacturer instructions, all buffers were supplemented with 60 U mL^−1^ Superase-In (Ambion) and 1× Halt Protease and Phosphatase Inhibitor Single-Use Cocktail (Life Technologies). Mitoplasts were obtained by incubating purified mitochondria in RNase A-containing hypotonic buffer (HEPES pH 7.2 supplemented with 1× Halt Protease and Phosphatase Inhibitor Single-Use Cocktail (Life Technologies) and 10 μg mL^−1^ RNase A (Roche)) for 20 min on ice and subsequently incubated for 10 additional min at room temperature in order to remove all possible cytosolic RNA contaminants. The purified mitoplasts were then washed thrice with Mitoplast Isolation Buffer (MIB: 250 nM Mannitol, 5 mM HEPES pH 7.2, 0.5 mM EGTA, 1 mg mL^−1^ BSA supplemented with 60 U mL^−1^ Superase-In (Ambion) and 1× Halt Protease and Phosphatase Inhibitor Single-Use Cocktail (Life Technologies)).

### Western blotting

Western blotting experiments were performed using the following primary antibodies: vinculin (V9131, clone hVIN-1 Sigma-Aldrich, 1:5,000), histone 3 (#4499, clone D1H2, Cell Signaling Technology, 1:1,000), MITF (ab12039, Abcam, 1:1,000), ATF4 (#11815, clone D4B8, Cell Signaling Technology, 1:1000), ATF5 (SAB4500895, Sigma-Aldrich, 1:500) eIF2α-Tot (#5324, clone D7D3, Cell Signaling Technology, 1:1000), eIF2α-Phospho-S51 (#3398, clone D9G8, Cell Signaling Technology, 1:1000), CHOP (#2895, clone L63F7, 1:1000), puromycin (MABE343, clone 12D10, Merck-Millipore, 1:10.000), β-actin (#4970, clone 13E5, Cell Signaling Technology, 1:1000), Tom20 (sc-17764, clone F-10, Santa Cruz Biotechnology, 1:1000). The following HRP-linked secondary antibodies were used: anti-mouse IgG (NA931-1ML, Sigma-Aldrich, 1:10,000) and anti-rabbit IgG (NA934-1ML, Sigma-Aldrich, 1:10,000). Relative protein levels were measured using ImageJ.

### PDX experiments

The cutaneous melanoma PDX models are part of the Trace collection (https://www.uzleuven-kuleuven.be/lki/trace/trace-leuven-pdx-platform) and were established using metastatic melanoma lesions derived from patients undergoing surgery as part of the standard treatment at the UZ Leuven. The uveal melanoma (UM) model derives from a lymph node metastasis and is a kind gift of M. Herlyn (The Winstar Institute, USA). Written informed consent was obtained from both patients and all procedures involving human samples were approved by the UZ Leuven/KU Leuven Medical Ethical Committee (S54185) and carried out in accordance with the principles of the Declaration of Helsinki. The experiments were approved by the KU Leuven animal ethical committee under ECD P038-2015 and performed in accordance with the internal, national and European guidelines of Animal Care and Use. Tumour pieces were implanted subcutaneously in the interscapular fat pad of female NMRI nude BomTac:NMRI-Foxn1^nu^, 4-week-old females (Taconic). Mice were maintained in a semi specific pathogen-free facility under standard housing conditions with continuous access to food and water. The health and welfare of the animals was supervised by a designated veterinarian. The KU Leuven animal facilities comply with all appropriate standards (cages, space per animal, temperature (22 °C), light, humidity, food, water), and all cages are enriched with materials that allow the animals to exert their natural behaviour. Mice used in the study were maintained on a diurnal 12-hour light/dark cycle. PDX models Mel-006, Mel-015, Mel-020 were derived from a female, male and female patients drug naïve patients respectively. The UM Mel-077 sample was derived from a male patient progressing on Pembrolizumab and Temozolomide. Mel-018, Mel-021 and Mel-078, are derived from male, female and male patients respectively. Once tumours reached 250 mm^3^ for Mel-077 and Mel-020, 500 mm^3^ for Mel-015 or 1000 mm^3^ for Mel-006, the mice were enrolled into treatment cohorts. Mice form distinct batches were randomly assigned to the different experimental groups. Mice were treated daily by oral gavage with a capped dose of 600-6 μg dabrafenib-trametinib respectively in 200 μL total volume, while Tigecycline (50 mg/kg) was administered daily by i.p. injection.

No specific randomization method was used. According to animal welfare guidelines, mice were sacrificed when tumours reach a volume of 2500 mm^3^ or when their body weight decreased more than 20% from the initial weight. Mice used in this paper never reached or overcame these limits.

### Allografts

1×10^5^ YUMM 1.7 and 5×10^4^ YUMMER 1.7 cells were injected subcutaneously in the interscapular fat pad with in C57BL/6 4-week-old males (males were chosen since females frequently rejected the implantation and had sporadic tumour ulceration). Mice were maintained under standard housing conditions with continuous access to food and water. The health and welfare of the animals was supervised by a designated veterinarian. The KU Leuven animal facilities comply with all appropriate standards (cages, space per animal, temperature (22 °C), light, humidity, food, water), and all cages are enriched with materials that allow the animals to exert their natural behaviour. Mice used in the study were maintained on a diurnal 12-hour light/dark cycle. The experiments were approved by the KU Leuven animal ethical committee under ECD P049-2019 and performed in accordance with the internal, national and European guidelines of Animal Care and Use. Once the tumours reached 100 mm^3^ (for YUMM 1.7) or 50 mm^3^ (for YUMMER 1.7), mice were enrolled in treatment cohort. Mice form distinct batches were randomly assigned to the different experimental groups. Mice were treated with Tigecycline (Bio Connect) i.p., (50 mg/kg) daily, with anti-PD-1 (Ultra-LEAF™ Purified anti-mouse CD279 antibody, clone RMP1-14) i.p., (10 mg/kg) twice per week for 3 weeks, for a total of 6 injections or a combination of the two (only for the YUMMER 1.7 allografts).

No specific randomization method was used. According to animal welfare guidelines, mice have to be sacrificed when tumours reach a volume of 2500 mm^3^ or when their body weight decreases more than 20% from the initial weight. Mice used in this paper were sacrificed upon reaching 1400 mm^3^ or, alternatively, upon reaching 16 (YUMM 1.7) or 19 (YUMMER 1.7) days of treatment.

### Immunofluorescence on PDX biopsies

Fluorescent staining was performed using OPAL staining reagents, which use individual tyramide signal amplification (TSA)-conjugated fluorophores to detect various targets within an immunofluorescence assay. In brief, samples were fixed with 4% Paraformaldehyde and embedded in paraffin. Serially cut sections of 5 μm were stained with haematoxylin and eosin for routine light microscopy, and used for immunohistochemistry.

Depending on the antibody, antigen retrieval was performed in Citrate buffer at pH 6 or EDTA buffer at pH 9. Deparaffinized sections were then incubated overnight with primary antibodies against AQP1 (cat No. #AB2219, Millipore, 1:3000), NGFR (cat No. 8238, Cell Signaling Technology, 1:1000), MITF (cat No. #HPA003259, Sigma-Aldrich, 1:100), MLANA (cat No. # HPA048662, Sigma-Aldrich, 1:200), CD36 (cat No. #HPA002018, Sigma-Aldrich, 1:200) and AXL (cat No. #AF154, R&D, 1:50), S100 (Cat#Z0311, Dako, 1:100). Subsequently, the slides were washed in phosphate-buffered saline, pH 7.2, and incubated for 10 min at room temperature with Opal Polymer HRP Mouse Plus Rabbit secondaries (PerkinElmer). After another wash in PBS, the slides were then incubated at RT for 10 min with one of the following Alexa Fluorescent tyramides (PerkinElmer) included in the Opal 4 color kit (Cat No. NEL810001KT, AKOYA Biosciences) to detect antibody staining, prepared according to the manufacturer’s instructions: Opal 520, Opal 570 and Opal 690 (dilution 1:50). Stripping of primary and secondary antibodies was performed by placing the slides in a plastic container filled with antigen retrieval (AR); microwave technology was used to bring the liquid at 100 °C (2 min), and the sections were then microwaved for an additional 15 min at 75 °C. Slides were allowed to cool in the AR buffer for 15 min at room temperature and were then rinsed with deionized water and 1 × Tris-buffered saline with Tween 20. After three additional washes in deionized water, the slides were counterstained with DAPI for 5 min and mounted with ProLong™ Gold Antifade Mountant (Cat No. P10144, ThermoFisher Scientific). Slides were scanned for image acquisition using Zeiss AxioScan Z.1 and ZEN2 software.

### Patient casistic

The patient was treated with BRAF-MEK inhibitors for stage IV malignant melanoma according to standard-of-care. PET-CT scans were done at baseline and for confirmation of remission. Every two months the patient had undergone radiographic assessment either with PET-CT or with a regular CT scan.

### Statistical analyses

All data are graphed as mean ± standard error of the mean (s.e.m.). In animal experiments, the designation ‘n’ indicates the number of animals used. When comparing two treatment cohorts over time, statistical significance was calculated by two-way ANOVA. When comparing 3 or more groups, statistical significance was calculated by one-way ANOVA with Dunnett’s multiple comparisons test. When comparing Kaplan-Meier-plots, statistical significance was calculated by Log-rank (Mantel-Cox) test. Statistical tests used are specified in the corresponding figure legends. Throughout all figures, statistical significance was concluded at *P*<0.05 (\**P*<0.05, \*\**P*<0.01, \*\*\**P*<0.001 and \*\*\*\**P*<0.0001). All statistical analyses were performed with GraphPad Prism v8.4.2 (464), April 08, 2020 for Mac OS Catalina.

