## Supplementary Table 1 and Supplementary Figures and Legends for "Activation of the Integrated Stress Response in drug-tolerant melanoma cells confers vulnerability to mitoribosome-targeting antibiotics"

| Primer | Sequence Forward (5' to 3') | Sequence Reverse (5' to 3') |
|---|---|---|
| MRPL53 | AACGTGGAATCGACGAGGAC | AGCATTTCCAGAGCGGTGAG |
| MRPL37 | GACCTGGACTGTAACGAGGG | GCAAACACATTTTGGTCCGC |
| MRPL40 | CATCAACACGCGCTATTGGG | CAAACGGCCCCGATCTTCTA |
| MRPL22P1 | CCACTGACCAGGCTTAGCTC | GGTCTTTCACTGCCACGTCT |
| ATP6VOB | GCTAGCACTGCTCTACTCCG | AACCATGCCACATCAAAGCG |
| ATP6VOA2 | GGTCAGCCTTACGAT GTCCA | GCAAACCCGGCTTTACGTTT |
| MRPL9 | TCCTCAGGGACTGGCTGTAT | ACAACACCAAGCGCCTCAC |
| 28S | TGTCGGCTCTTCCTACCATTGT | ACCCAGCTCACGTTCCCTATTA |
| 18S | TTCGGAACTGAGGCCATGAT | TTTCGCTCTGGTCCGTCTTG |

**Supplementary Table legend**

**Supplementary Table 1 | qPCR primers used in the study.**

**Supplementary Figure Legends**

**Figure S1 | Tigecycline reduces melanoma cell growth *in vitro*. A,** Cell growth (measured as % of cell confluency) of YUMM 1.7 (resistant to immunotherapy) and YUMMER 1. (immunotherapy-sensitive) cell lines upon exposure to increasing concentrations of Tigecycline for 72 hours. Data are the means ± s.e.m. of three independent experiments. *P* values were calculated by Dunnett's test. **B,** (Left) Colony formation assays with cells described in in **Figure 2A** and in **A** treated with increasing concentrations of Tigecycline (Tige). The violet colour is due to crystal violet, a compound that binds intracellular DNA and protein, thus highlighting the cells attached to the plate. Representative image of three independent experiments. (Right) Quantification of colony formation assays of cells described in **Figure 2A** and in **A** presented as the mean density (percentage of area occupancy) ± s.e.m. of three independent experiments. *P* values were calculated by Dunnett's test. $^{**}P<0.01$, $^{***}P<0.001$, $^{****}P<0.0001$.

**Figure S2 | Tigecycline increases overall survival *in vivo*. A,** Kaplan-Meier-plot showing overall survival of mice described in **Figure 2C**. *P* value was calculated by log-rank (Mantel-Cox) test. **B,** Kaplan-Meier-plot showing overall survival of Mel-006 $BRAF^{V600E}$ PDX mice treated with vehicle (DMSO) or Tigecycline. *P* value was calculated by log-rank (Mantel-Cox) test. **C,** Kaplan-Meier-plot showing overall survival of Mel-015 $BRAF^{V600E}$ PDX mice treated with vehicle (DMSO) or Tigecycline. *P* value was calculated by log-rank (Mantel-Cox) test. **D,** Kaplan-Meier-plot showing overall survival of YUMM 1.7 mouse xenografts treated with $\alpha$-PD-1 or untreated (Ctrl). *P* value was calculated by log-rank (Mantel-Cox) test. **E,** Tumour volume of YUMMER 1.7 mouse xenografts described in **Figure 2F**. *NS* = $P>0.5$, $^{*}P<0.05$.

**Figure S3 | Tetracyclines affect the growth of multiple drug-tolerant states. A,** (Left) Colony formation assays with cells described in in **Figure 3A**. The violet colour is due to crystal violet, a compound that binds intracellular DNA and protein, thus highlighting the cells attached to the plate. Representative image of three independent experiments. (Right) Quantification of colony formation assays of cells described in **Figure 3A** presented as the mean density (percentage of area occupancy) ± s.e.m. of three independent experiments. **B,** Caspase activity (measured as average # of caspase$^{+}$ cells per image) of cells described in **Figurs 2B and 3B**. Data are presented as the mean ± s.e.m. of three independent experiments *P* values were calculated by Dunnett's test. *P* values were calculated by Dunnett's test. $^{*}P<0.05$, $^{**}P<0.01$, $^{****}P<0.0001$.

**Figure S4 | Tigecycline significantly increases overall survival. A,** Kaplan-Meier-plot showing overll survival of mice described in **Figure 4A**. *P* values were calculated by log-rank (Mantel-Cox) test. **B,** Kaplan-Meier-plot showing overall survival of mice described in **Figure 4C**. *P* values were calculated by log-rank (Mantel-Cox) test. *NS:* $P>0.05$, $^{**}P<0.01$, $^{***}P<0.001$, $^{****}P<0.0001$.

**Figure S5 | Activation of the ISR predicts durable responses to antibiotic treatment. A,** Western blotting of Mel-020 (*NRAS*mut) PDX tumours collected right before the start of the treatment ($T_0$) or at the end of the experiment ($T_{end}$) after receiving a daily dose of vehicle or Tigecycline. **B,** Western blotting of Mel-077 (Uveal melanoma) PDX tumours collected right before the start of the treatment ($T_0$) or at the end of the experiment ($T_{end}$) after receiving a daily dose of vehicle or Tigecycline. **C,**

Western blotting of cells described in **Figure 6C**. Representative image of three independent experiments.

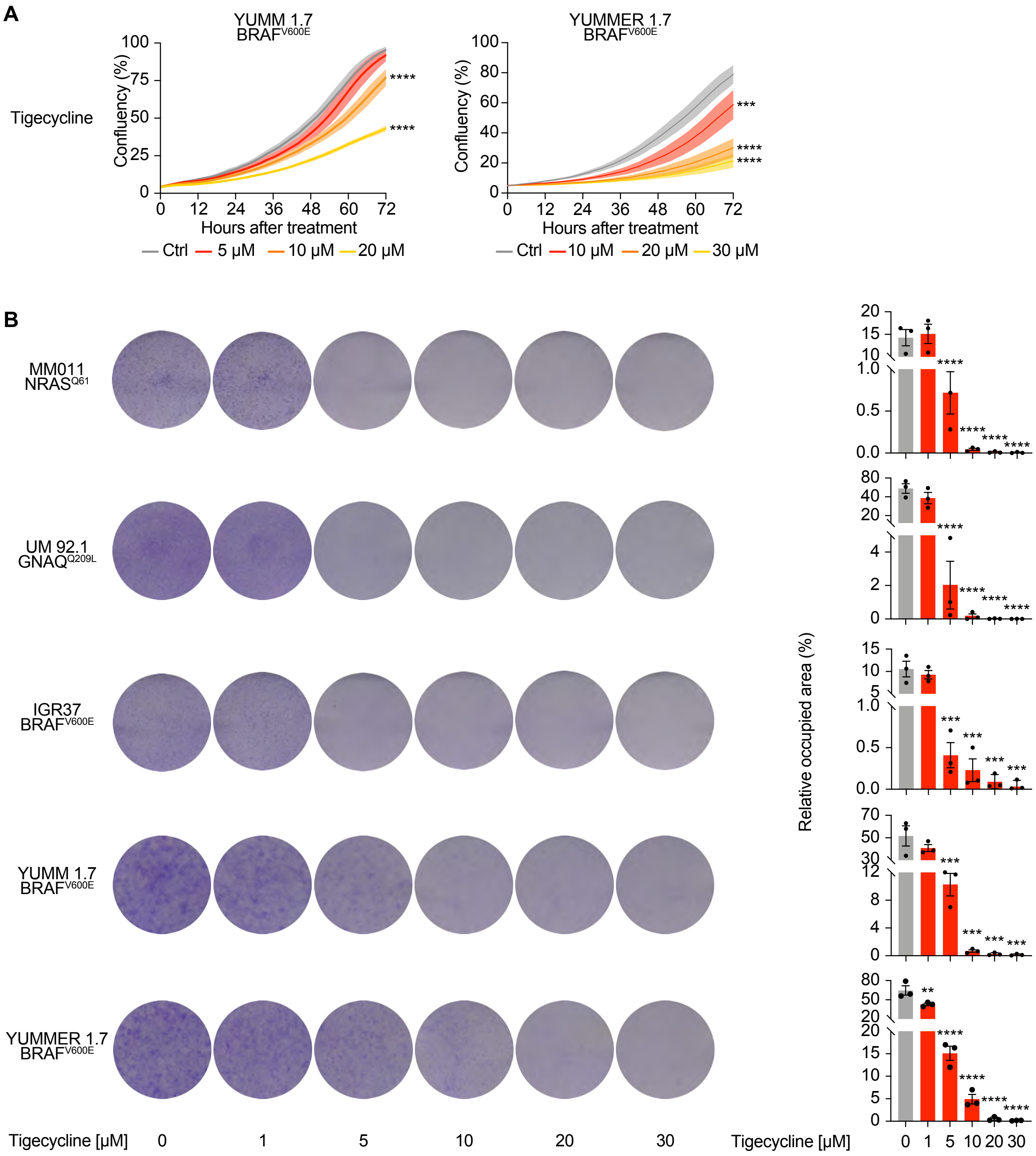


**Figure S1 | Tigecycline reduces melanoma cell growth *in vitro*.**

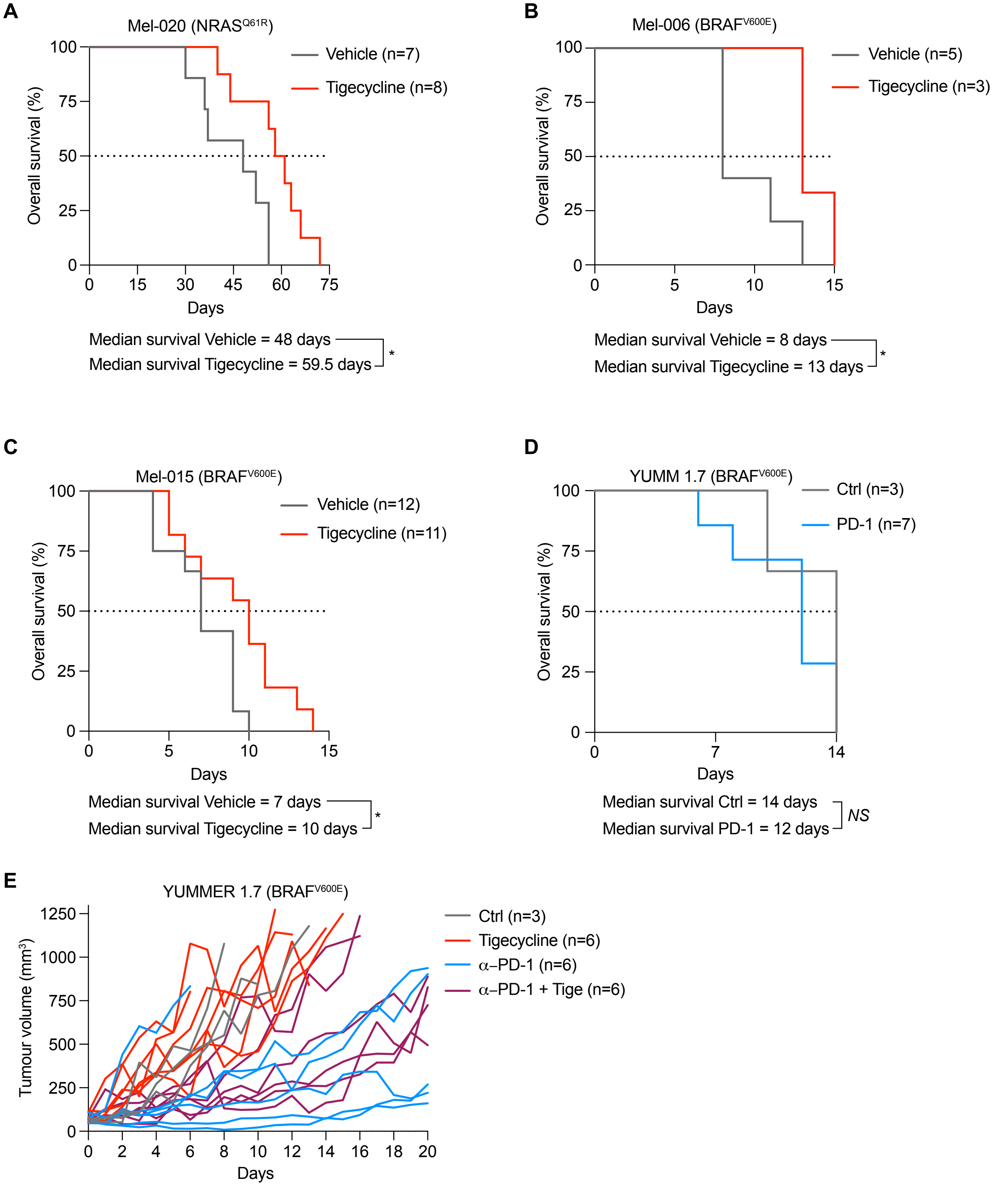


**Figure S2 | Tigecycline increases overall survival *in vivo*.**

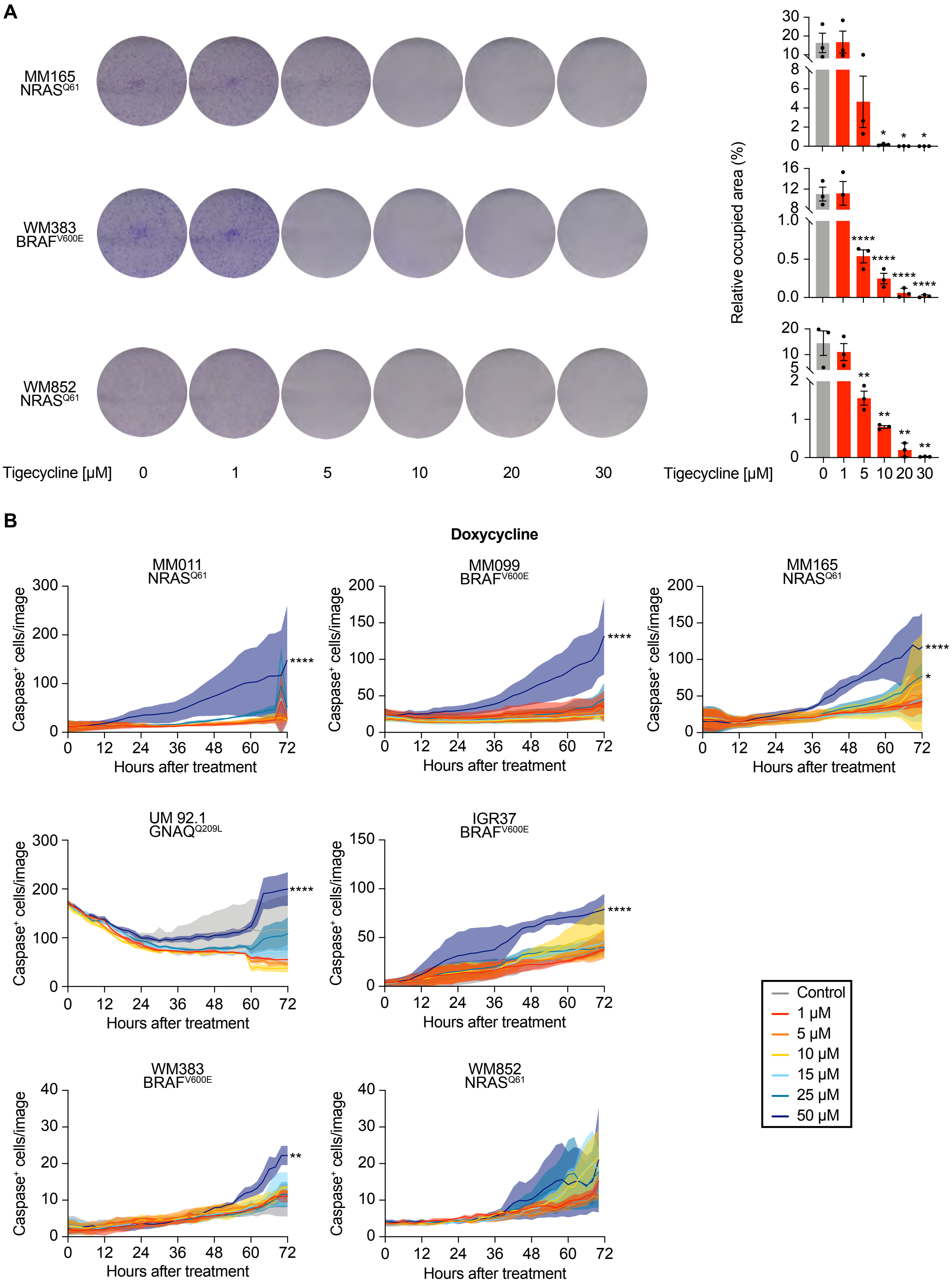


**Figure S3 | Tetracyclines affect the growth of multiple drug-tolerant states.**

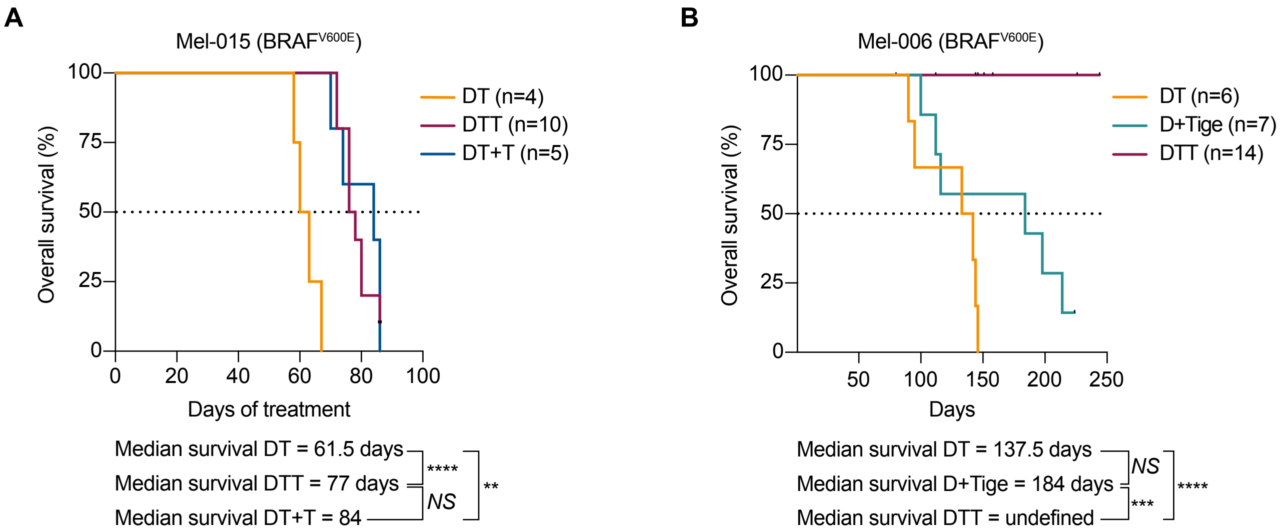


**Figure S4 | Tigecycline significantly increases overall survival.**

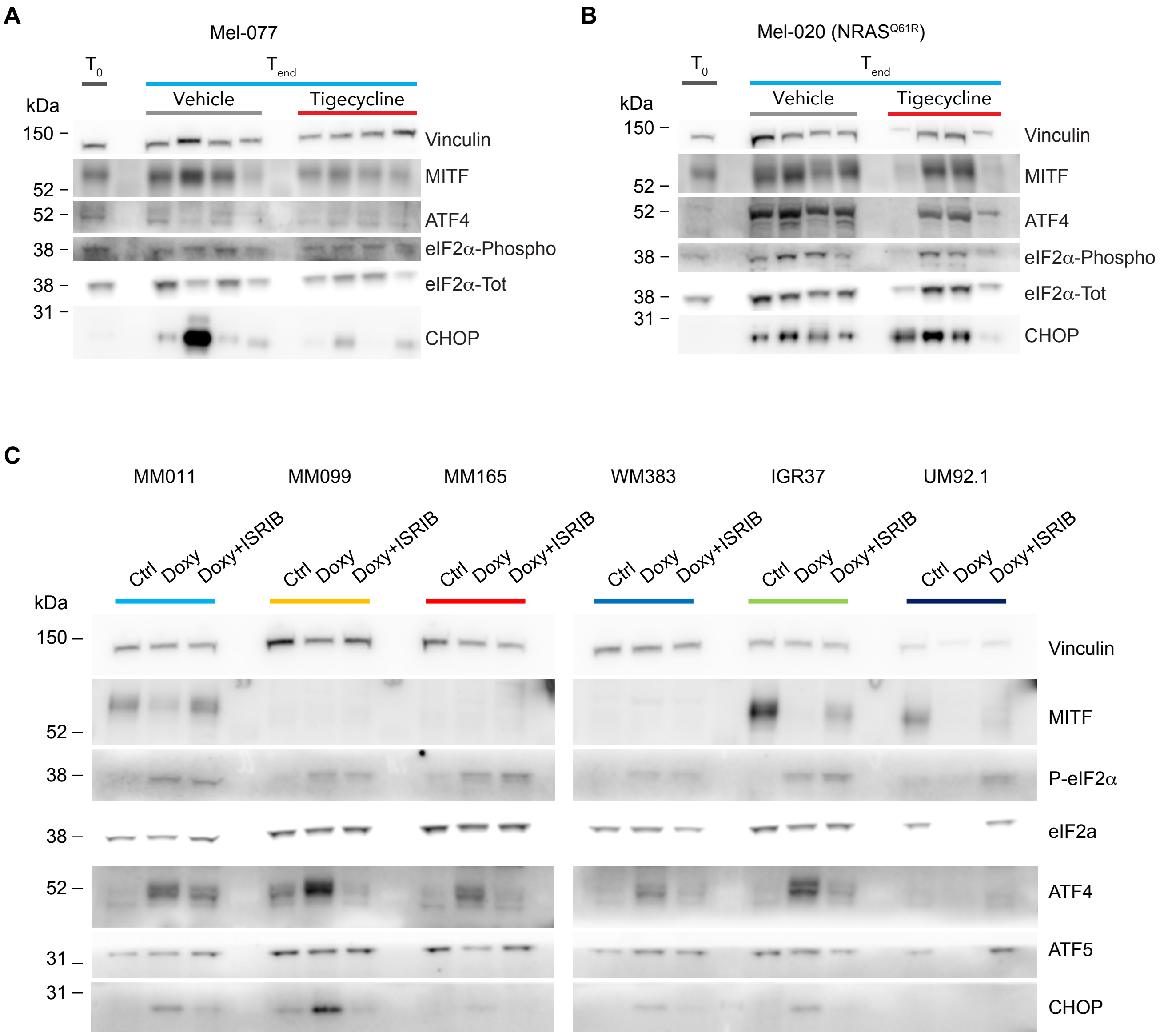


**Figure S5 | Activation of the ISR predicts durable responses to antibiotic treatment.**
